# Evolution of neocortical folding: A phylogenetic comparative analysis of MRI from 34 primate species

**DOI:** 10.1101/379750

**Authors:** Katja Heuer, Omer Faruk Gulban, Pierre-Louis Bazin, Anastasia Osoianu, Romain Valabregue, Mathieu Santin, Marc Herbin, Roberto Toro

## Abstract

We conducted a comparative analysis of primate cerebral size and neocortical folding using magnetic resonance imaging data from 65 individuals belonging to 34 different species. We measured several neocortical folding parameters and studied their evolution using phylogenetic comparative methods. Our results suggest that the most likely model for neuroanatomical evolution is one where differences appear randomly (the Brownian Motion model), however, alternative models cannot be completely ruled out. We present estimations of the ancestral primate phenotypes as well as estimations of the rates of phenotypic change. Based on the Brownian Motion model, the common ancestor of primates may have had a folded cerebrum similar to that of a small lemur such as the aye-aye. Finally, we observed a non-linear relationship between fold wavelength and fold depth with cerebral volume. In particular, gyrencephalic primate neocortices across different groups exhibited a strikingly stable fold wavelength of about 12 mm (± 20%), despite a 20-fold variation in cerebral volume. We discuss our results in the context of current theories of neocortical folding.

## Introduction

The human brain is the largest and most folded of those of extant primates. Much discussion has surrounded the question of whether its characteristics are due to a specific selection or to random drift. On the one hand, the large human brain may be just an expected result of descent with modification: That a primate brain has the volume of ours could not be surprising, as it would not be surprising to throw 10 times heads if we toss a coin a large enough number of times. On the other hand, a large and profusely folded brain could be a selected trait, providing a significant adaptive advantage – the substrate for the sophisticated cognitive abilities that have enabled humans to thrive, multiply, and invest almost all ecosystems on earth.

The evolution of human neuroanatomy has been studied for many years, with contradictory results regarding the question of a human exception. While several studies have suggested that different human neuroanatomical traits are outside the general primate trend (Rilling and Insel 1999, Schoenemann et al 2005, Gazzaniga 2008), many others see a continuation (Prothero and Sundsten 1984, Zilles et al 1989, Semendeferi et al 2002, Herculano-Houzel 2009). A potential issue of most of these studies was the lack of an appropriate account of phylogenetic relationships. Phylogenetic relationships introduce violations of the assumption of statistical independence of observations: the phenotypes of closely related species are expected to be more similar than those of distant species. Indeed, even 2 completely random variables can appear as correlated if they are allowed to vary along a phylogenetic tree (Felsenstein 1985).

Phylogenetic comparative methods aim at using information on the development and diversification of species – phylogenies – to test evolutionary hypotheses (Nunn and Barton 2001, Nunn 2011). Today, gene sequencing allows us to build phylogenetic trees based on the differences across homologous genes in various species. It is furthermore possible to use the number of changes necessary to match the gene sequences of one species into those of another to estimate their time of divergence from a hypothetical common ancestor (Paradis 2012). The lengths of the phylogenetic tree branches can then be made to represent the time of the progressive splits of the species at the tips of the tree, from a series of common ancestors (the nodes of the tree).

Given such phylogenetic trees, we can build and test models of the evolution of phenotypic traits under different hypotheses. Three influential models of trait evolution are the Brownian Motion model (BM), the Ornstein-Uhlenbeck model (OU), and the Early-Burst model (EB). The Brownian Motion model supposes that phenotypic changes diffuse randomly along the tree (Cavalli-Sforza and Edwards 1967, Felsenstein 1973, 1985). The phenotype of two species having split early from their common ancestor will then be less similar than that of species having recently split. The Ornstein-Uhlenbeck model supposes that phenotypic changes are not completely random, but tend towards specific values (Lande 1976, Hansen 1997, Cooper et al 2015). These could be values which are particularly advantageous and have therefore a higher probability of being selected. Finally, the Early-Burst model (Harmon et al 2010) considers the possibility that phenotypic changes are initially faster (when a new adaptive regime is first invested), and then slow-down.

Several recent studies have adopted phylogenetic comparative analysis methods to study the evolution of primate neuroanatomy (Smaers et al 2011, Barton and Venditti 2013a, Lewitus et al 2014, Miller et al 2019). In particular, a series of reports have considered the question of the exceptionality of the size of the human prefrontal cortex relative to other primate species (Smaers et al 2011, 2017, Barton and Venditti 2013a, b). Some of these reports suggest an exceptionally large and significantly more asymmetric prefrontal cortex (Smaers et al 2011, 2017, Smaers 2013), whereas others find it to be as large as expected (Barton and Venditti 2013a, b, Miller et al 2019). The problem does not appear to be settled, but the availability of published data on prefrontal grey and white matter volume has allowed researchers to contrast their different methodological approaches using the same data.

Here we present a phylogenetic comparative analysis of primate neuroanatomy based on a sample of magnetic resonance imaging (MRI) data from 65 specimens coming from 34 different primate species. We acquired and made openly available high-resolution MRI data for 33 specimens from 31 different species. This is part of an ongoing effort to digitise the Vertebrate Brain Collection of the National Natural History Museum of Paris. The remaining 33 specimens come from different openly accessible sources. All the data has been indexed in the collaborative neuroimaging website BrainBox (http://brainbox.pasteur.fr, Heuer et al. 2016), to facilitate access and foster community-driven data analysis projects. This dataset can be used to perform detailed analyses of neocortical anatomy, beyond volumetric measurements. We looked at several global measurements of neocortical folding, including estimations of global gyrification, total folding length, average fold wavelength and average fold depth. After considering various alternative evolutionary models, our results indicate that the BM model provided the best fit to the data, suggesting that random change may be a main force in primate neuroanatomical evolution. Based on the BM model, we provide estimations of the ancestral values for the different phenotypes under study, as well as estimations of the evolutionary rates of phenotypic change. All our analyses scripts have been made available in an accompanying GitHub repository: https://github.com/neuroanatomy/34primates.

## Methods

### Data Sources

Magnetic resonance imaging (MRI) data was obtained for 66 individuals across 34 different primate species. Thirty one brains from 29 species were obtained from the Vertebrate Brain Collection of the National Museum of Natural History (MNHN) of Paris (see information on Data Acquisition below). Eleven MRI datasets were downloaded from our Brain Catalogue website (https://braincatalogue.org): one crab-eating macaque, one gorilla, and 9 chimpanzees donated by the National Chimpanzee Brain Resource (NCBR, kindly provided by Chet Sherwood and William D. Hopkins, http://www.chimpanzeebrain.org). The bonobo, gibbon and a second gorilla were downloaded from NCBR, from within the data provided by James Rilling and Thomas Insel (Rilling and Insel, 1999). Three further macaque datasets, one rhesus macaque and two crab-eating macaques, were kindly provided by the Pruszynski Lab and downloaded from Zenodo (Arbuckle et al 2018). 8 additional macaque datasets, 4 rhesus and 4 crab-eating macaques, were downloaded from the IoN site of PrimeDE (Milham et al 2018). Finally, the surfaces from 10 human brains were selected and downloaded from the New York site of the ABIDE 1 dataset, through the ABIDE preprocessed project (http://preprocessed-connectomes-project.org/abide, Craddock et al 2013). These subjects were unaffected controls, 20 to 30 years old, and had a good quality surface reconstruction upon expert visual examination. A list of the included species can be found in Table 1. Scripts to automatically download these datasets are available in the accompanying GitHub repository: https://github.com/neuroanatomy/34primates.

**Table 1.** List of species included. ABIDE 1: Autism Brain Imaging Data Exchange 1. BC: Brain Catalogue. MNHN: Muséum Nationale d’Histoire Naturelle de Paris. NCBR: National Chimpanzee Brain Resource. PL: Pruszynski Lab. PDE: PRIMate Data Exchange (PRIME-DE).

| Name | Binomial Name (GenBank) | N | In vivo | Extracted | Provenance |
| --- | --- | --- | --- | --- | --- |
| **Lemuriformes** |  |  |  |  |  |
| Aye-aye | Daubentonia madagascariensis | 1 | No | No | MNHN |
| Black-and-white ruffed lemur | Varecia variegata variegata | 1 | No | No | MNHN |
| Coquerel's mouse lemur | Mirza coquereli | 1 | No | No | MNHN |
| Grey mouse lemur | Microcebus murinus | 1 | No | No | MNHN |
| Mongoose lemur | Eulemur mongoz | 1 | No | No | MNHN |
| Red-tailed sportive lemur | Lepilemur ruficaudatus | 1 | No | No | MNHN |
| Ring-tailed lemur | Lemur catta | 1 | No | Yes | MNHN |
| **Loridae** |  |  |  |  |  |
| Red slender loris | Loris tardigradus | 1 | No | Yes | MNHN |
| **Galagonidae** |  |  |  |  |  |
| Demidoff's galago | Galago demidoff | 1 | No | No | MNHN |
| **Cebidae** |  |  |  |  |  |
| Black-pencilled marmoset | Callithrix penicillata | 1 | No | Yes | MNHN |
| Cotton-top tamarin | Saguinus oedipus | 1 | No | Yes | MNHN |
| Douroucouli | Aotus trivirgatus | 1 | No | No | MNHN |
| Squirrel monkey | Saimiri sciureus | 2 | No | Yes | MNHN |
| Tufted capuchin | Cebus apella | 1 | No | No | MNHN |
| White-faced sapajou | Cebus capucinus | 1 | No | Yes | MNHN |
| **Atelidae** |  |  |  |  |  |
| Black spider monkey | Ateles paniscus | 2 | No | No | MNHN |
| Woolly monkey | Lagothrix lagotricha | 1 | No | Yes | MNHN |
| **Cercopithecini** |  |  |  |  |  |
| Green monkey | Chlorocebus sabaeus | 1 | No | Yes | MNHN |
| Moustached guenon | Cercopithecus cephus cephus | 1 | No | Yes | MNHN |
| **Papionini** |  |  |  |  |  |
| Crab-eating macaque | Macaca fascicularis | 8 | No, Yes, Yes | Yes, No, No | BC, PL, PDE |
| Grey-cheeked mangabey | Lophocebus albigena | 1 | No | Yes | MNHN |
| Hamadryas baboon | Papio hamadryas | 1 | No | Yes | MNHN |
| Rhesus monkey | Macaca mulatta | 6 | No, Yes, Yes | Yes, No, No | MNHN, PL, PDE |
| Sooty mangabey | Cercocebus atys | 1 | No | Yes | MNHN |
| **Colobinae** |  |  |  |  |  |
| Hanuman langur | Semnopithecus entellus | 1 | No | Yes | MNHN |
| Indochinese lutung | Trachypithecus germaini | 1 | No | No | MNHN |
| King colobus | Colobus polykomos | 1 | No | Yes | MNHN |
| **Hominoidea** |  |  |  |  |  |
| Bonobo | Pan paniscus | 1 | Yes | No | NCBR |
| Chimpanzee | Pan troglodytes troglodytes | 9 | Yes | No | NCBR |
| Gibbon | Hylobates lar | 1 | Yes | No | NCBR |
| Gorilla | Gorilla beringei | 1 | No | Yes | BC |
| Gorilla | Gorilla gorilla | 1 | Yes | No | NCBR |
| Human | Homo sapiens | 10 | Yes | No | ABIDE 1 |
| Orangutan | Pongo pygmaeus | 1 | No | No | MNHN |

### Data Acquisition

The 31 brains from the Vertebrate Brain Collection of the MNHN were scanned at the Center for Neuroimaging Research (CENIR) of the Institut du Cerveau et de la Moëlle Épinière (ICM, Paris, France). High resolution MRI images were acquired using either a 3T Siemens Tim Trio system, a 3T Siemens Prisma, or an 11.7T Bruker Biospec. Each dataset was acquired with a 3D gradient-echo sequence (FLASH). Parameters (Field of View, Matrix size, TR, TE) were adjusted so as to obtain the highest resolution possible with our scanner (from 100 to 450 µm isotropic). TR and TE were always chosen as minimum. Flip angle was fixed to 20° at 3T and 10° at 11.7T. The number of averages was chosen to maintain a scanning time below 12 hours.

### Data Preprocessing

The MRI data from the MNHN was converted to Nifti 1 format (Cox et al 2004) using FLS 5.0.10 (Jenkinson et al 2012, https://fsl.fmrib.ox.ac.uk) and dcm2niix (Chris Rorden, version v1.0.20170724, https://www.nitrc.org/projects/mricrogl/). We used our web tool Reorient (https://github.com/neuroanatomy/reorient) to rotate the brains so that the sagittal plane was always straight, the superior/inferior directions were respected (we cannot verify whether the left/right orientations are correct, we only assume they are, and we visually check that no flips were introduced by our analysis pipeline), and the axis of the corpus callosum is horizontal. We also used Reorient to crop the brains. All this MRI data was uploaded to Zenodo (https://zenodo.org), and the links are provided in the accompanying GitHub repository.

The chimpanzees, the bonobo, the gibbon and the gorilla from the NCBR were converted to Nifti 1 format, and we used the DenoiseImage tool included in ANTs (Avants et al 2009) to improve the signal-to-noise ratio. The data was then reoriented and cropped using Reorient, contrast inhomogeneities were corrected using N4BiasFieldCorrection from ANTS (Tustison et al 2010), and finally the intensity range was manually limited to prevent regions with high intensity from affecting the global contrast (usually the optic nerves). The chimpanzee data was processed using Freesurfer (Dale et al 1999, Fischl et al 2001) using the script recon-all-chimps.sh in the accompanying GitHub repository.

### Data quality control

Our data comprise post mortem as well as in vivo MRI scans and vary in tissue conservation, resolution and signal-to-noise ratio. Some of the MRIs include only the extracted brain, others include the brain and the skull, and finally a few others include the entire body of the animal. We generated images of one coronal, axial and sagittal slice using Nilearn (Abraham et al 2014) to perform a first visual quality control. A thorough visual quality control was later performed during the manual segmentation stage.

Quantitative indications of data quality were obtained by measuring signal-to-noise ratios: We computed the ratio of the signal of interest divided by the standard deviation of the background (region without signal). The background region of interest was automatically detected by identifying the first peak in the MRI’s histogram.

The signal of interest was defined based on the histogram of the MRI after removing the voxels from the background. We then detected the histogram’s peak and selected the position of maximum density. When several peaks were detected, we excluded the first one – most often related to CSF or the fixative fluid in ex vivo brains – and used the average position of the remaining peaks as the mean signal of interest. The results are provided in Supplemental Table S1.

### Manual segmentation and surface reconstruction

All the MRI data, except for the chimpanzees and the humans, were segmented using BrainBox (Heuer et al 2016, http://brainbox.pasteur.fr), a Web application for the visualisation, annotation and real-time collaborative segmentation of MRI data. Offline, we used StereotaxicRAMON, Thresholdmann, Segmentator and ITK-SNAP to generate a first mask of the cerebrum. StereotaxicRAMON (https://github.com/neuroanatomy/StereotaxicRAMON) provides manual editing tools, a series of topology-preserving mathematical morphology operators, as well as a real-time 3D visualisation of the manual segmentations. Thresholdmann (https://github.com/neuroanatomy/thresholdmann) enables the generation of binary segmentation masks by using a threshold that can be adjusted locally: the value of the threshold at intermediate points is then interpolated using radial basis functions. Segmentator (Gulban et al 2018a, b, https://github.com/ofgulban/segmentator) enables the generation of binary segmentation masks through the interactive manipulation of a 2-D histogram where the x-axis represents grey level and the y-axis represents the magnitude of the gradient at each point of the image. Finally, ITK-SNAP (Yushkevich et al 2006, http://www.itksnap.org) is a general tool for manual medical image segmentation.

The semi-automatically obtained masks were uploaded to BrainBox, where we created a project centralising all the data. The BrainBox project can be accessed here: http://brainbox.pasteur.fr/project/BrainCataloguePrimates. The main manual segmentation tasks performed in BrainBox involved removal of the cerebellum, brainstem and optic nerves; delineation of sulci missed by the automatic segmentation; and reconstruction of damaged tissue parts (see Figure 1 for examples). After manual segmentation was finished and reviewed by at least one more person, we implemented a script in Python 3.6 to download all the data using BrainBox’s RESTful API (the script is included in the accompanying GitHub repository). The manually segmented masks were then transformed into triangular meshes using the CBS tools (Bazin et al 2014, https://www.nitrc.org/projects/cbs-tools). The workflow included the following steps: mask binarisation, transformation of the mask into a probability function, and extraction of an isosurface using the connectivity-consistent Marching Cubes algorithm (Han et al 2003).

**Figure 1.**
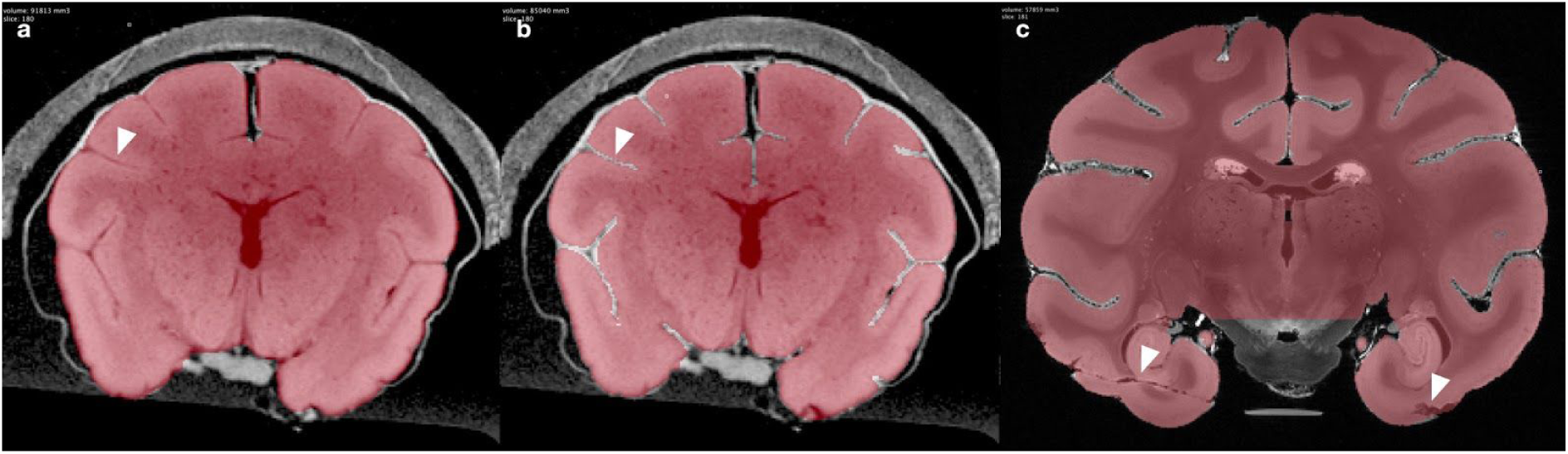
Examples of segmentation tasks. (a) Cerebrum masks were obtained using different semi-automatic methods. These masks often failed to properly segment sulci, as pointed by the white arrow. (b) Manual segmentation of the cerebrum involved the removal of the cerebellum, brainstem and optic nerves, and the exclusion of the sulci, (c) as well as reconstructing damaged tissue parts as pointed by the arrow.

All meshes were then processed using the following steps: soft Laplacian smoothing to remove the shape of voxels, decimation down to 3 vertices per mm^2^ (https://github.com/cnr-isti-vclab/vcglib/tree/master/apps/tridecimator), removal of isolated vertices, and non-shrinking Taubin smoothing (Taubin 1995), implemented in our tool Mesh Geometry (https://github.com/neuroanatomy/meshgeometry), to remove further geometric artefacts.

### Neuroanatomical measurements

We used Mesh Geometry to compute the volume, surface, absolute gyrification index, folding length, and estimated number of sulci for each mesh. We define the absolute gyrification index as the ratio between the surface of a cerebral hemisphere mesh and the surface of a sphere of the same volume as the hemisphere’s volume. Because the sphere is the solid with the least surface for a given volume, this provides an absolute index of the “excess” of surface of a cerebrum: A sphere has then an absolute gyrification index of 1 (the minimum), and in our measurements a human cerebrum has an absolute gyrification index of about 4. The folding length measures the total length of the curves dividing sulci from gyri (as measured using a mean curvature map). This measurement is conceptually similar to the gyral length measurement used by Prothero and Sundsten (1984) or the sulcal lengths referenced by Zilles et al (1989), however, those measurements were performed only on the surface or even in endocasts. The estimated number of sulci is obtained by counting all the regions with negative mean curvature (the sulci).

We use the cerebrum mesh surface area (*S*), volume (*V*) and the folding length (*L*) to estimate the average wavelength (*W*) and depth of the folds (*D*) in a cerebrum (see Figure 2). The total surface of a cerebrum mesh can be thought as the multiplication of its total folding length times the average profile of a fold (the curve that goes from an inflexion point, up to the gyral crest, down to the sulcal fundus, and up to the next inflexion point). Furthermore, we can use the convex hull of each hemisphere – scaled to have the same volume *V* as the hemisphere – to estimate the total surface of its hypothetical unfolded version. In this unfolded version of the hemisphere the profile of a fold is simply the wavelength of the fold (as the sulcal depth is 0). The average wavelength (*W*) of folding in the cerebrum can be estimated as the ratio between the surface area of the convex hull (*S_h_*) and half the total folding length (*L*):

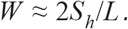

**Figure 2.**
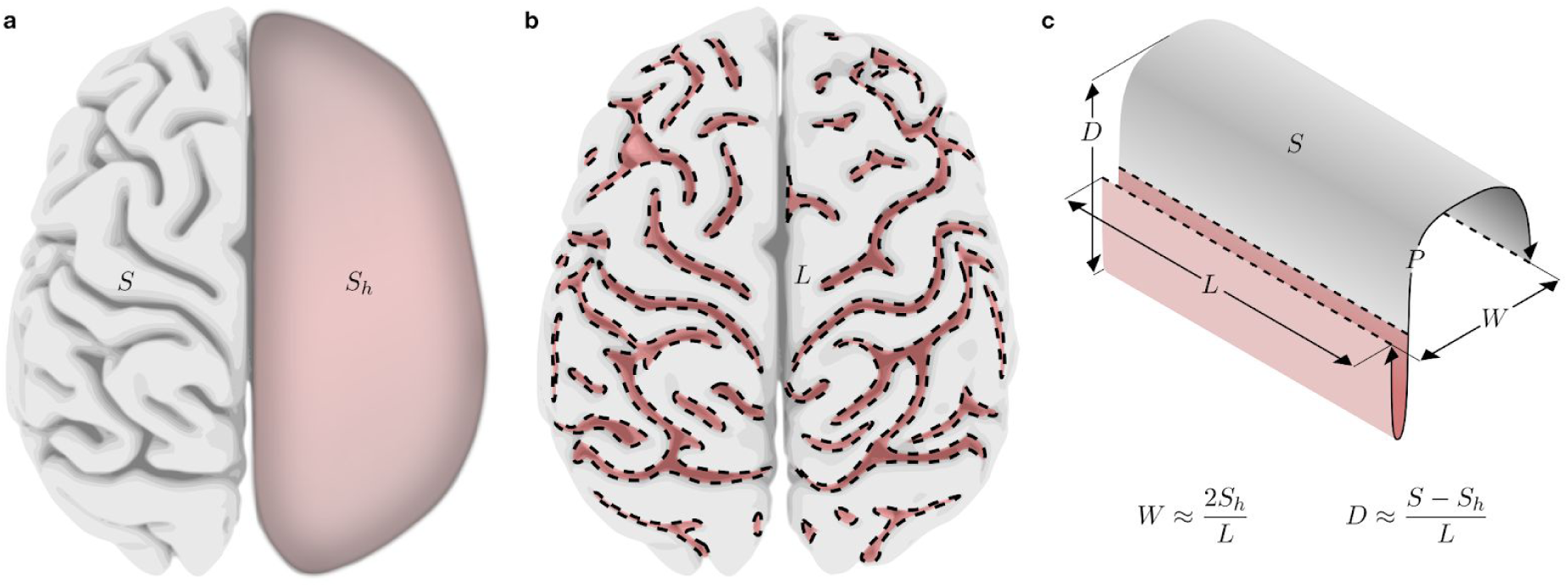
Neuroanatomical measurements. (a) Absolute gyrification index as the “excess” of the cerebral surface over the surface of its convex hull, normalised to have the same volume. Here illustrated over the surface of a bonobo brain. (b) Folding length – the total length of the curves dividing sulci from gyri, shown as dashed lines over the surface of a bonobo brain. (c) Schematic illustration of the neuroanatomical measurements used in our equations.

If we further approximate, as Prothero and Sundsten (1984), the profile of a fold to be like a square function (folds going straight up and down), we have that the total surface of the cerebrum mesh should be:

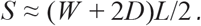

In the case of the hypothetical unfolded version of the mesh, the surface should be

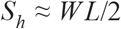

(because D=0). We then have that

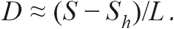

We used Mesh Geometry to split the left and right hemispheres, and the command line qhull (Barber et al 1996, http://www.qhull.org) to compute their convex hulls. The scripts necessary to compute all these measurements are available in the accompanying GitHub repository.

### Statistics and phylogenetic comparative analyses

We downloaded phylogenetic tree data for our 34 primate species from the 10k trees website (Arnold et al 2010, https://10ktrees.nunn-lab.org/Primates/downloadTrees.php). This website provides a Bayesian inference of primate phylogeny based on 17 genes. We obtained the consensus tree as well as a sample of 100 trees in proportion to their posterior probabilities.

We used R 3.5.0 (R Core Team 2018, http://www.R-project.org) for our statistical analysis. Measurements of surface area, volume, folding length and folding number varied over several orders of magnitude and were log-transformed before analysis. Phylogenetic independent contrasts (PICs, Felsenstein 1985) were computed using the packages ape (Paradis 2012) and phytools (Revell 2011), with multiple observations per species.

We fitted different evolutionary models (Brownian Motion, Ornstein-Uhlenbeck with a single alpha, with one alpha per phenotype, with a full multivariate matrix of alpha values, and the Early Burst model) using the package Rphylopars (Goolsby et al. 2016), which allows the analysis of multivariate phenotypes with multiple observations per species.

The Brownian Motion (BM) model supposes that phenotypes diffuse randomly through the branches of the phylogenetic tree with intensity controlled by the parameter σ (σ^2^ is the variance of the Brownian process). Under the BM model, phenotypes of species that have diverged recently should then be more similar than those of species that have diverged earlier. The Ornstein-Uhlenbeck (OU) model supposes that phenotypic variation is not only random, but is also attracted to an evolutionarily advantageous value, with a strength controlled by the parametre alpha (when alpha=0, the OU model is equivalent to the BM model). Finally, the Early Burst (EB, Harmon et al 2010) model supposes that the speed of phenotypic change can be faster at some point (when a new adaptive zone is invested) and slow-down after. When the rate parametre r in the EB model is r=0, the model reduces to the BM model, and negative values indicate rates of change that decrease through time.

We used the Akaike Information Criterion (AIC) values for the fit of each of these models to select among them (a smaller value indicates a better fit). Following the criteria of Burnham and Anderson (2004), we considered that an AIC difference between 4 to 7 suggests considerable less support for the model with larger AIC value, and a difference larger than 10 suggests essentially no support for the model with larger AIC value.

## Results

### Data collected

We obtained cerebrum surface reconstructions for 65 individuals from 34 different primate species (we excluded only 1 specimen from the original 66 datasets collected, a red howler monkey, due to extensive tissue damage. The MRI is nevertheless available in the BrainBox project as well as in Supplemental Table S1). Table 1 displays the complete list of species included, the number of individuals per species, and information on provenance. Figure 3 shows dorsal views of our reconstructions conserving a homogeneous scale (only one individual per species). The amount of neocortical folding was strongly related to cerebral volume: small *Strepsirrhini* primates had a basically unfolded neocortex, as well as several of the *Platyrrhini* primates (New World monkeys) in our sample. When folds appeared, their pattern was strongly left-right symmetric in small cerebra and became progressively more asymmetric in the larger *Papionini* and *Hominoidea* brains.

**Figure 3.**
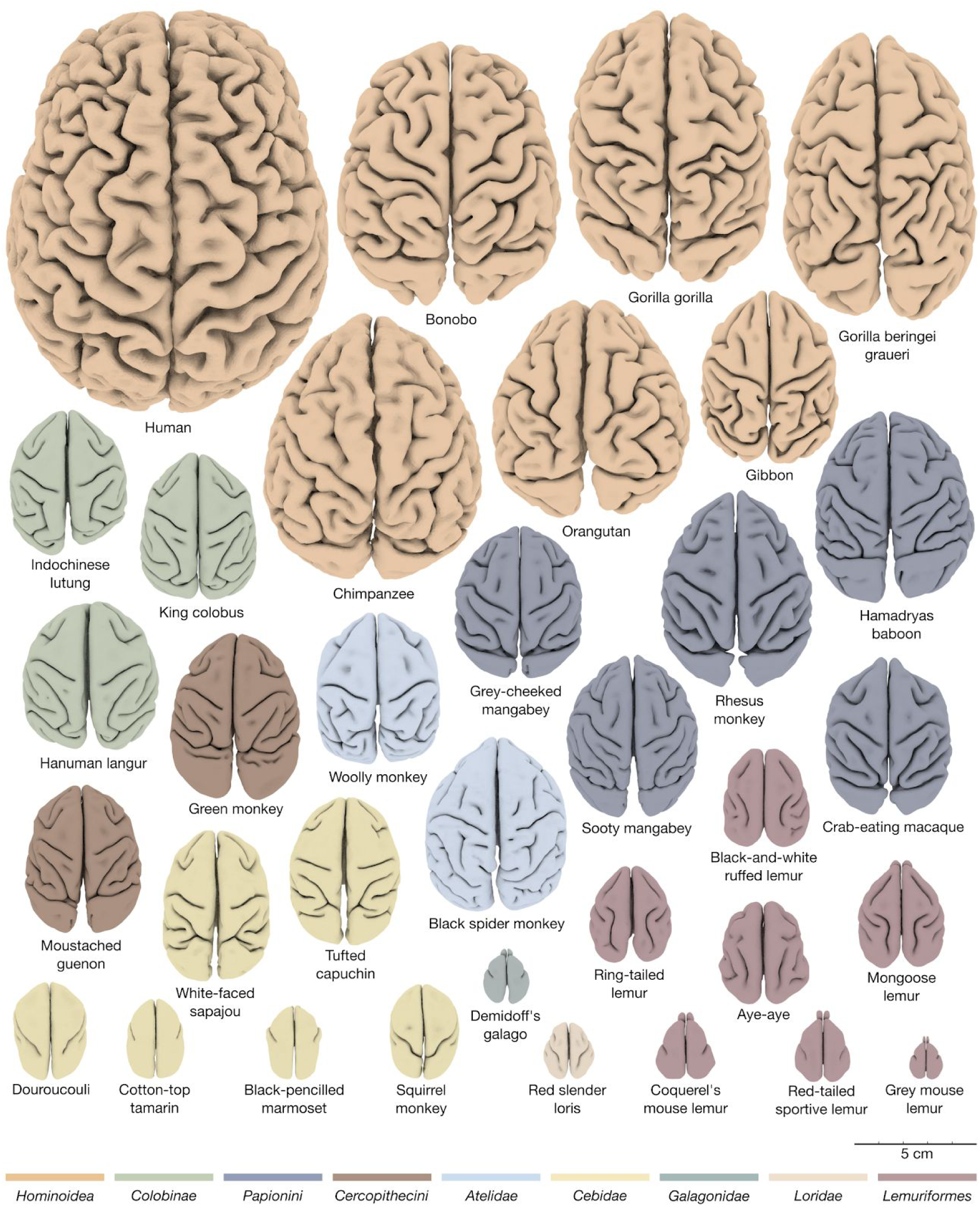
Dorsal view of the reconstructed cerebral hemispheres of 34 different primate species. Colours represent the different clades, and brains are represented from largest on top to smallest at the bottom. The scale is the same for all brains. High-res version: https://doi.org/10.5281/zenodo.2538751

### Neuroanatomical measurements

The relationships among all our neuroanatomical measurements are illustrated in Figure 4. Surface area and volume correlated strongly (R^2^=0.99, p≪1) with a positive allometric scaling coefficient beta=0.82. There was also a strong positive correlation with our absolute gyrification index (AbsGI), total folding length, and folding number count. Our estimations of average fold wavelength and average fold depth exhibited an interesting, non-linear relationship with cerebral volume. Despite a > 3-fold variation in volume between humans and chimpanzees, and a > 20-fold variation in volume between humans and the crab-eating macaque, the average fold wavelength changed only from about 11 mm in the human sample, 12 mm in the chimpanzee sample and the bonobo (less than 1.1-fold), to 14 mm in the crab-eating macaque sample (less than 1.3-fold). In the group of primates with small cerebra, the estimation of average fold wavelength is to be interpreted cautiously. It was often the case that a few folds would develop in a largely smooth cerebrum, rendering the notion of wavelength difficult. This can be observed in Figure 5a, which shows the relationship between cerebral volume and average fold wavelength. What could be interpreted as very wide folds in the smaller cerebra may reflect indeed the presence of a single fold within an essentially lissencephalic cerebrum. Interestingly, as cerebral volume increases and the notion of wavelength becomes more relevant, we observe a progressive stabilisation of the fold wavelength. A similar but opposite trend can be observed for our estimation of the average fold depth (Figure 5b). For small, lissencephalic cerebra, the value is close to 0 (as expected), increases rapidly with cerebral volume up to 6 mm, and tends to stabilise and increase slowly up to 10 mm in humans (8 mm in the chimpanzees and the bonobo, 6 mm in the crab-eating macaque, Figure 5b).

**Figure 4.**
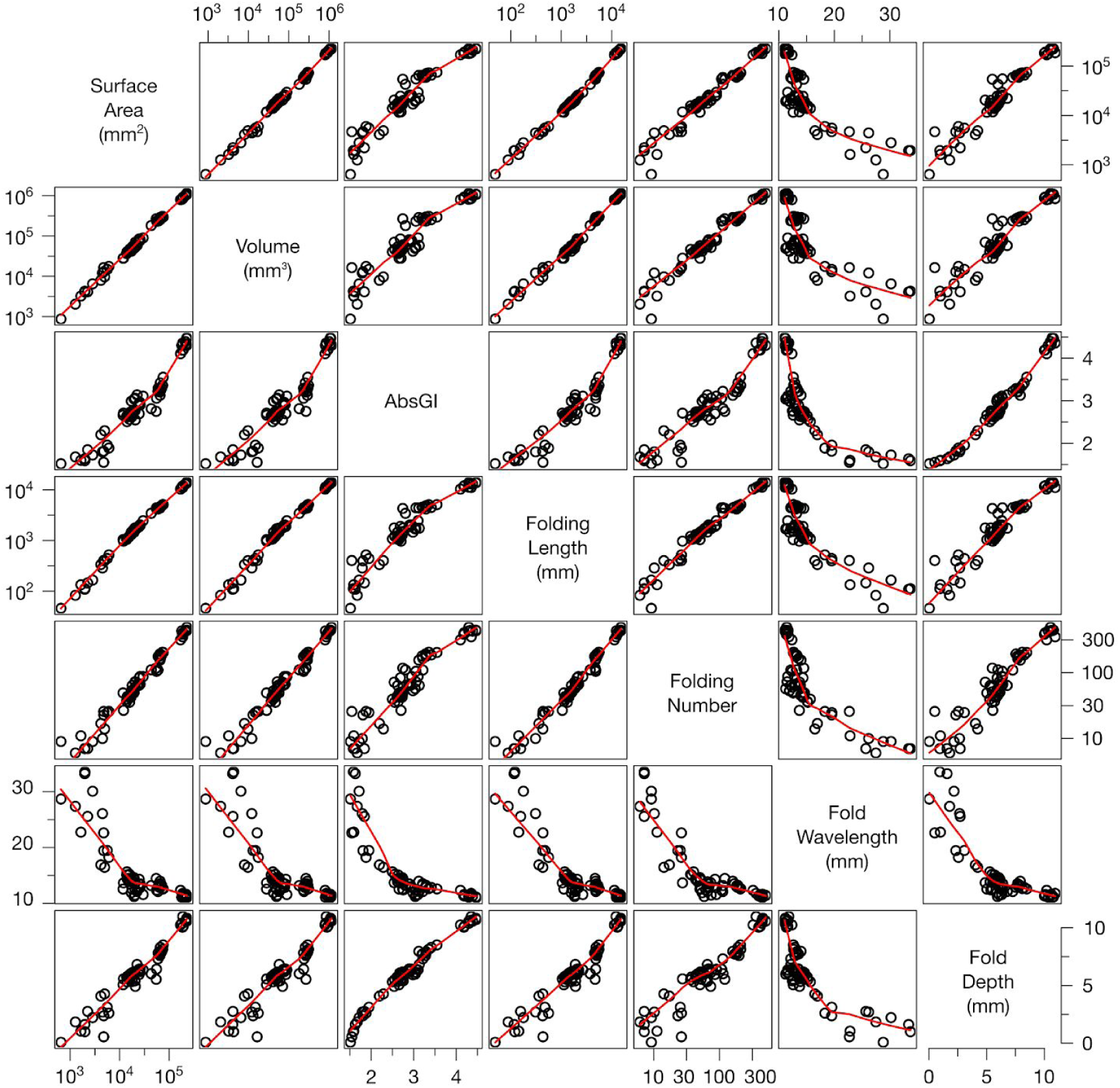
Scatterplots of neuroanatomical measurements. The scatterplots show the correlation between all pairs of measurements used in this study. A Log10 transformation was used on measurements that varied over several orders of magnitude, such as surface area or volume. The red curve is a locally estimated scatterplot smoothing (LOESS).

**Figure 5.**
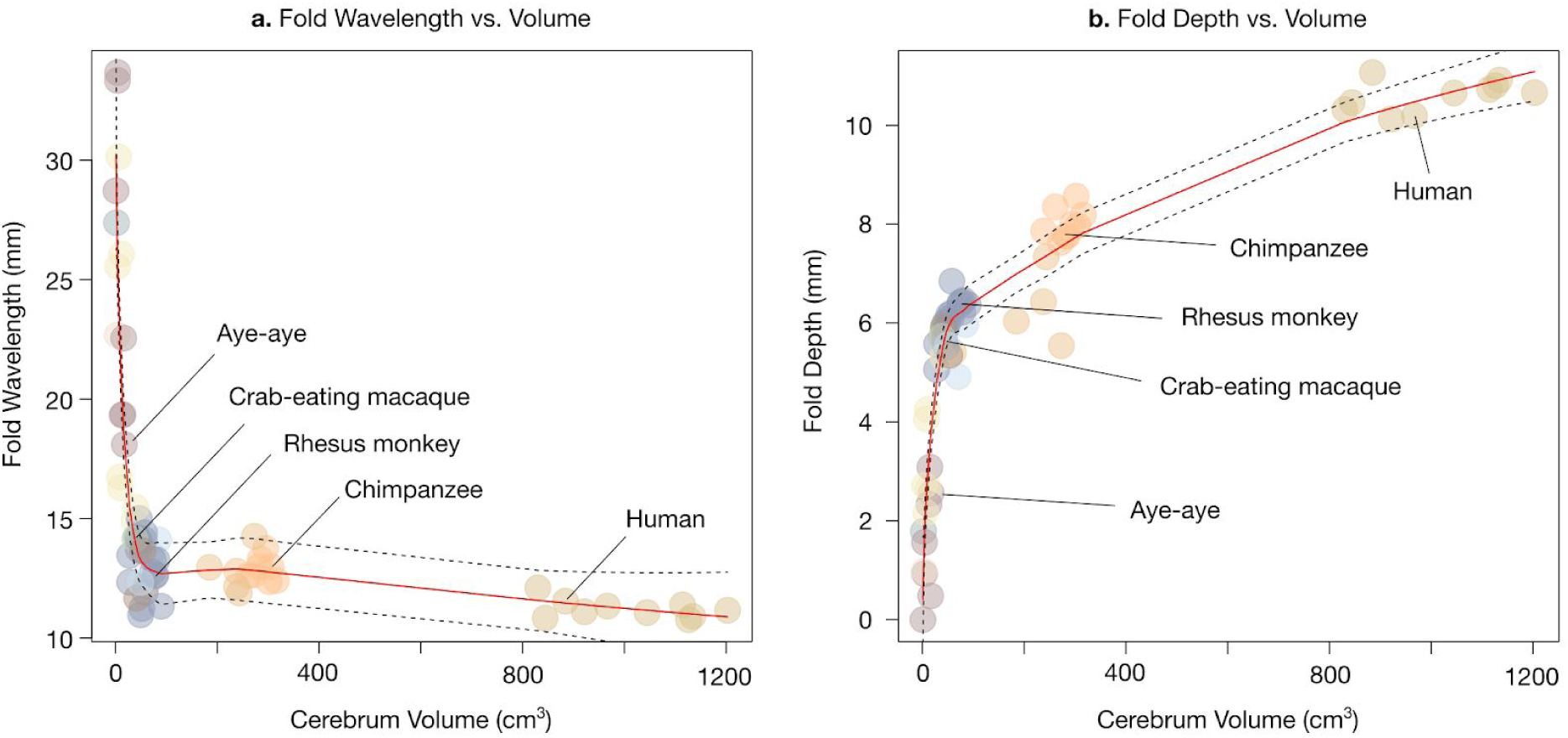
Relationship between cerebral volume, fold wavelength and fold depth. (a) Fold wavelength versus cerebral volume. Fold wavelength was conserved among almost ⅔ of species with relatively large brains. Among the remaining species with small, lissencephalic, brains the estimation of fold wavelength is not well defined and provides values which roughly correspond to the size of the brain. (b) Fold depth versus cerebral volume. The relationship between fold depth and cerebral volume also shows an inflexion point separating species with small and large brains. It increases steeply when brains have only a few folds (which become deep rapidly) and more softly when brains have profuse folding. The colours of the data points correspond to their clades, and several representative species are annotated to facilitate comparison.

### Phylogenetic comparative neuroanatomical analyses

Figure 6 shows the consensus phylogenetic tree used in our analyses. The branch length represents an estimation of time since split from a common ancestor. The number of specimens per species is indicated in parenthesis, and we provide a grouping of different species (tips of the tree) in families and clades. The best fit for the variation of neuroanatomical phenotypes along the phylogenetic tree was obtained for the BM model: a random change in phenotypes with variability depending on phylogenetic distance. The differences in model fit (AIC values) suggest considerably less support for the 2^nd^ and 3^rd^ best models – the OU model with a single alpha value, and the EB model – and essentially no support for all the other models (see Table 2). Our following analyses focus therefore on the results obtained assuming the BM model.

**Table 2.** Phylogenetic model selection. Different models of phenotypic evolution were fitted to the data and ranked by their AIC (smaller values indicate a better fit).

| Model | AIC |
| --- | --- |
| Brownian Motion (Pagel's $\lambda=1$) | -956.31 |
| Ornstein-Uhlenbeck, single alpha | -948.97 |
| Early Burst | -947.10 |
| Star (Pagel's $\lambda=0$) | -923.41 |
| Ornstein-Uhlenbeck, diagonal alpha matrix | -909.79 |
| Ornstein-Uhlenbeck, full alpha matrix | -698.53 |

**Figure 6.**
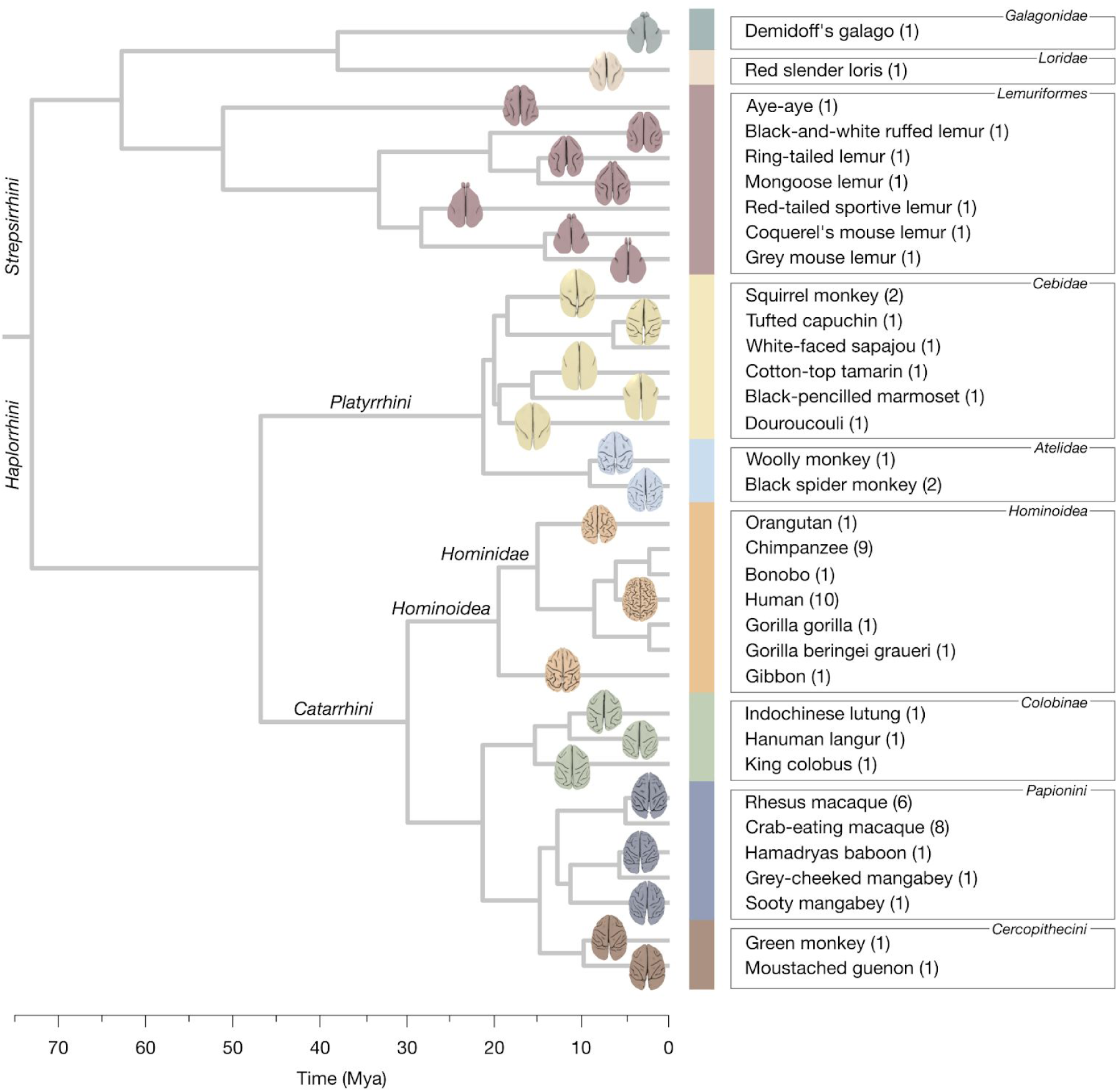
Phylogenetic tree. The phylogenetic tree represents a Bayesian inference of primate phylogeny based on genotyping data of 17 genes. The time of split of tree branches is provided in a scale of millions of years ago (Mya). The colour bar as well as the colours of the brains represent their clades (as in Fig. 3). The number of MRIs used for each species is provided in parenthesis besides each species’ name.

The analyses of phenotypic relationships including phylogenetic information (Figure 7) agree in essence with the previous analyses of the raw data (Figure 4). The role of phylogeny is, however, strong and statistically significant. Pagel (1999) suggested a strategy to test for the importance of the phylogenetic signal which relies on a modification of the branch lengths (and therefore of the phylogenetic variance-covariance matrix). The out-of-diagonal elements of the variance-covariance matrix are multiplied by a value λ, 0≤λ≤1. When λ=1, the results are equivalent to those of the BM model. When λ=0, all species are supposed to be independent (producing a phylogenetic tree with a “star” shape). The log-likelihood of the λ=1 model was strongly significantly larger than that of the λ=0 model (534.3 versus 517.7, χ^2^=33.1, p-value≪1).

**Figure 7.**
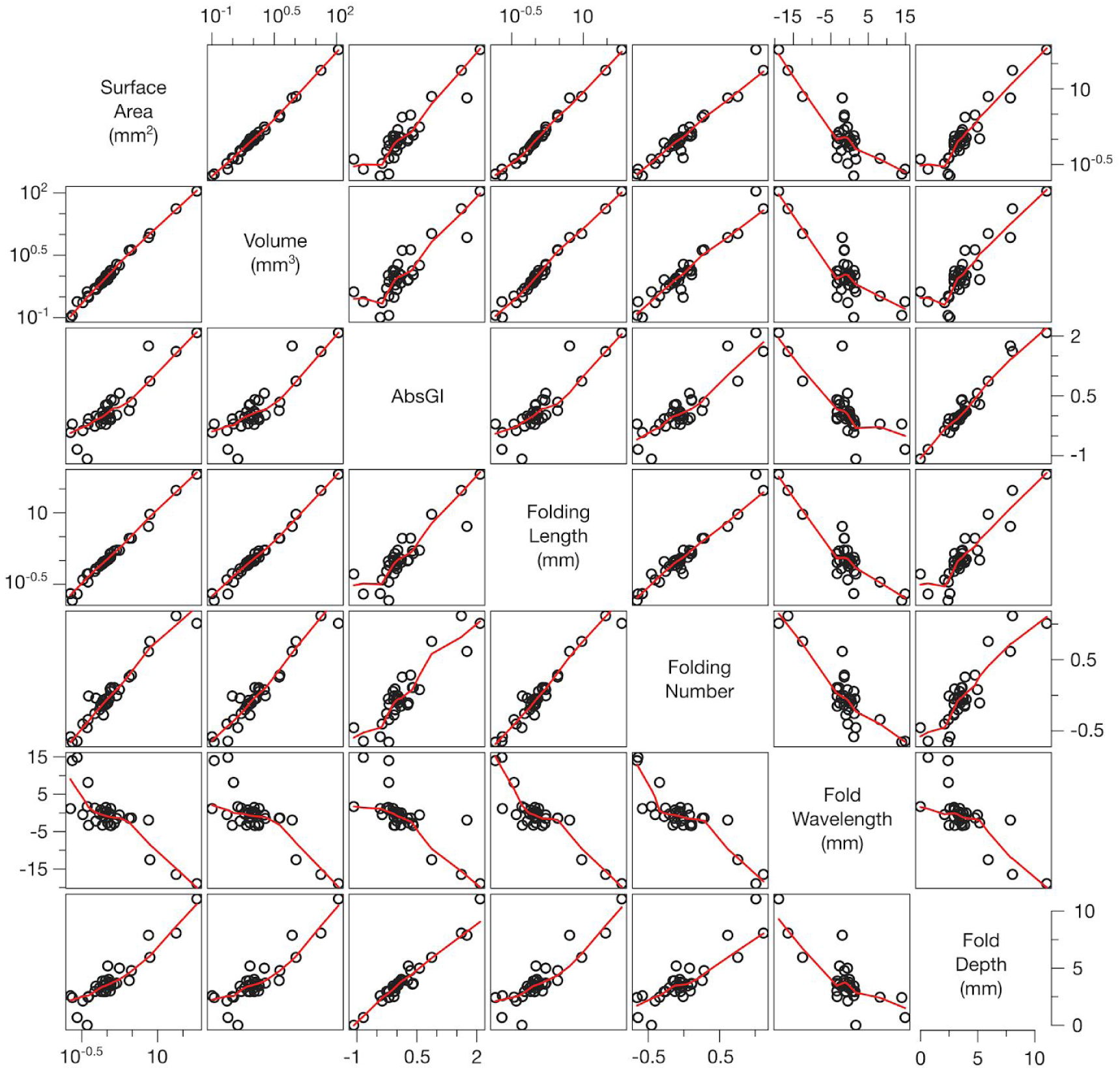
Phylogenetic comparisons of the neuroanatomical phenotypes. Scatterplots comparing each pair of neuroanatomical measurements, taking into account the phylogenetic relationships. The consensus phylogenetic tree was used to obtain phylogenetic independent contrasts (PIC), which were then used in the comparisons. Measurements varying over several orders of magnitude were Log10 converted.

The estimation of ancestral neuroanatomical phenotypes based on the BM model suggests a general increase in cerebral volume and neocortical surface in the *Catarrhini* branch, progressing along the *Hominoidea* branch and reaching its maximum among *Homininis* (Figure 8). Interestingly, within the *Platyrrhini* branch, both within the *Cebidae* family (the tufted capuchin) and the *Atelidae* family, some species exhibit an increase in cerebral volume, which corresponds with an increase in the number of folds and the emergence of neocortical folding asymmetries. The phenograms (Figure 9) show a different perspective on the same data, where the vertical axis represents time, the horizontal axis represents phenotype, and the phylogenetic relationships are represented by a branching pattern linking the phenotypes of extant species with those predicted for their common ancestors. We can observe a continuous evolutionary increase in cerebral volume, surface area, folding length, etc., from the common ancestor of all primates to humans (highlighted in red), but more complex patterns of increases and decreases for other species. Interestingly, we can also see that for the largest part of species with a highly folded neocortex, the average folding wavelength clusters in a small range between 11 to 14 mm (highlighted in blue).

**Figure 8.**
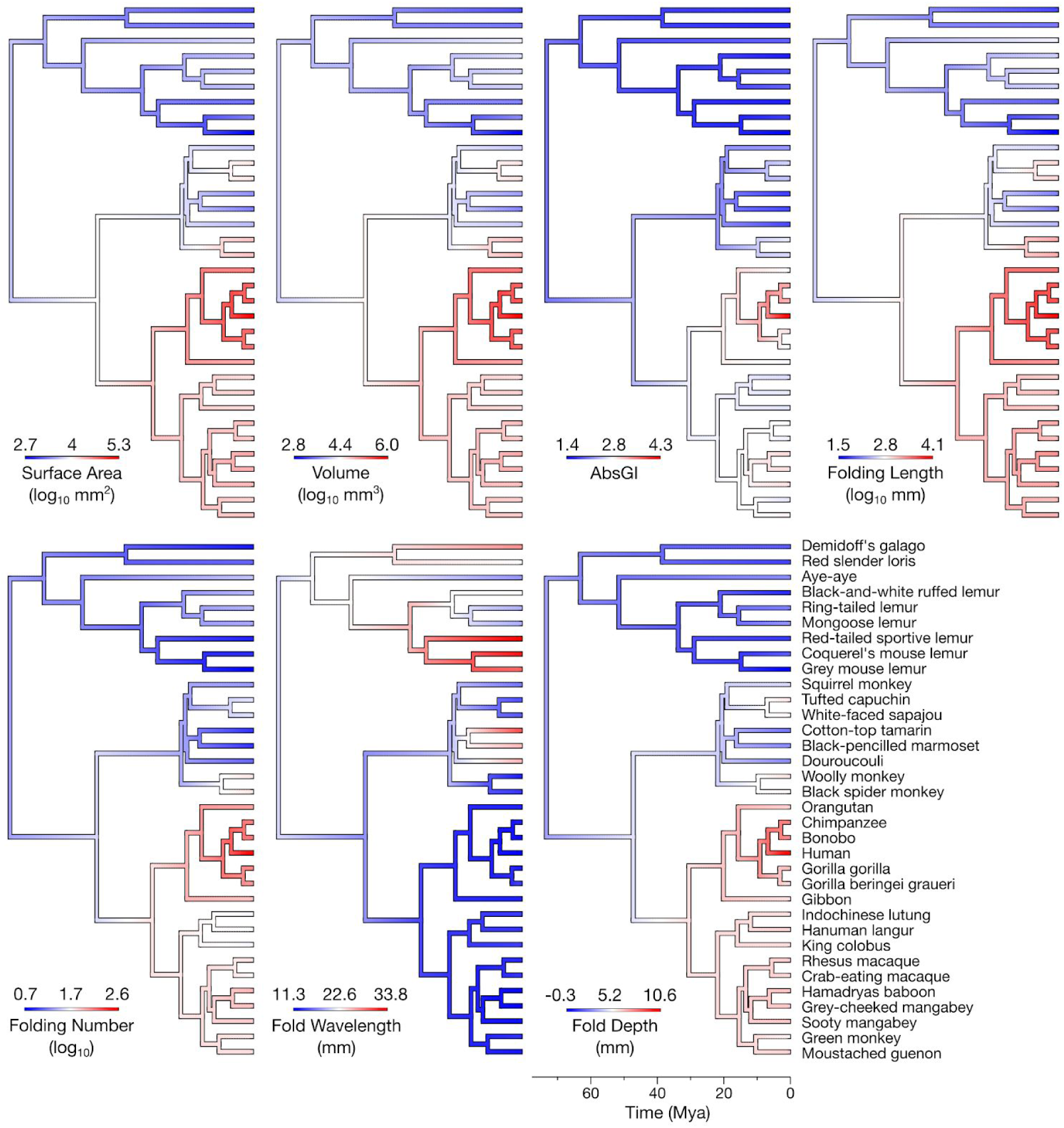
Estimated ancestral neuroanatomical phenotypes. The ancestral estimations of each phenotype were obtained using a Brownian Motion model of phenotypic evolution. Their values are represented in colour over the consensus phylogenetic tree. The species at the tip of the tree are indicated in the lower-right tree.

**Figure 9.**
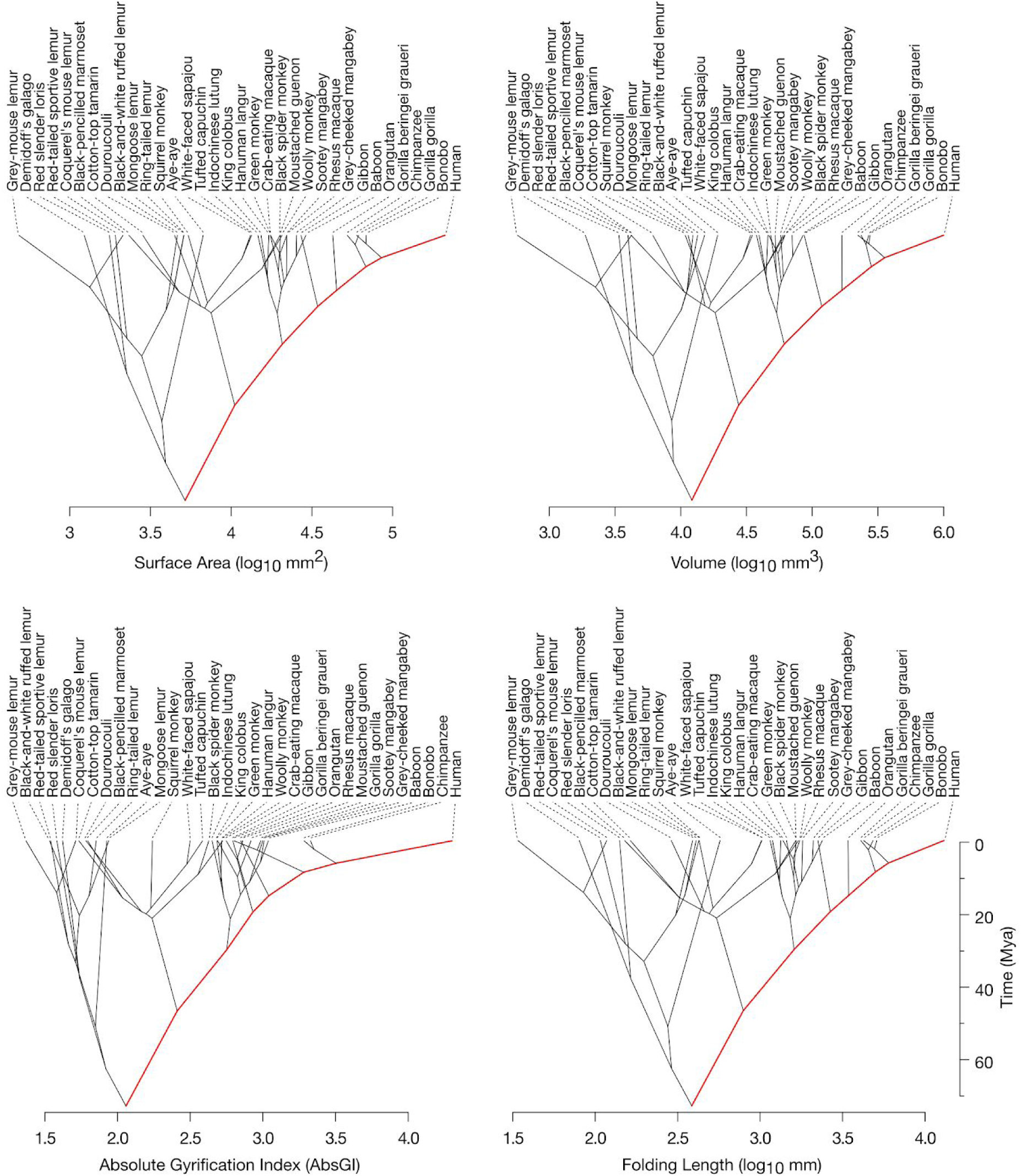
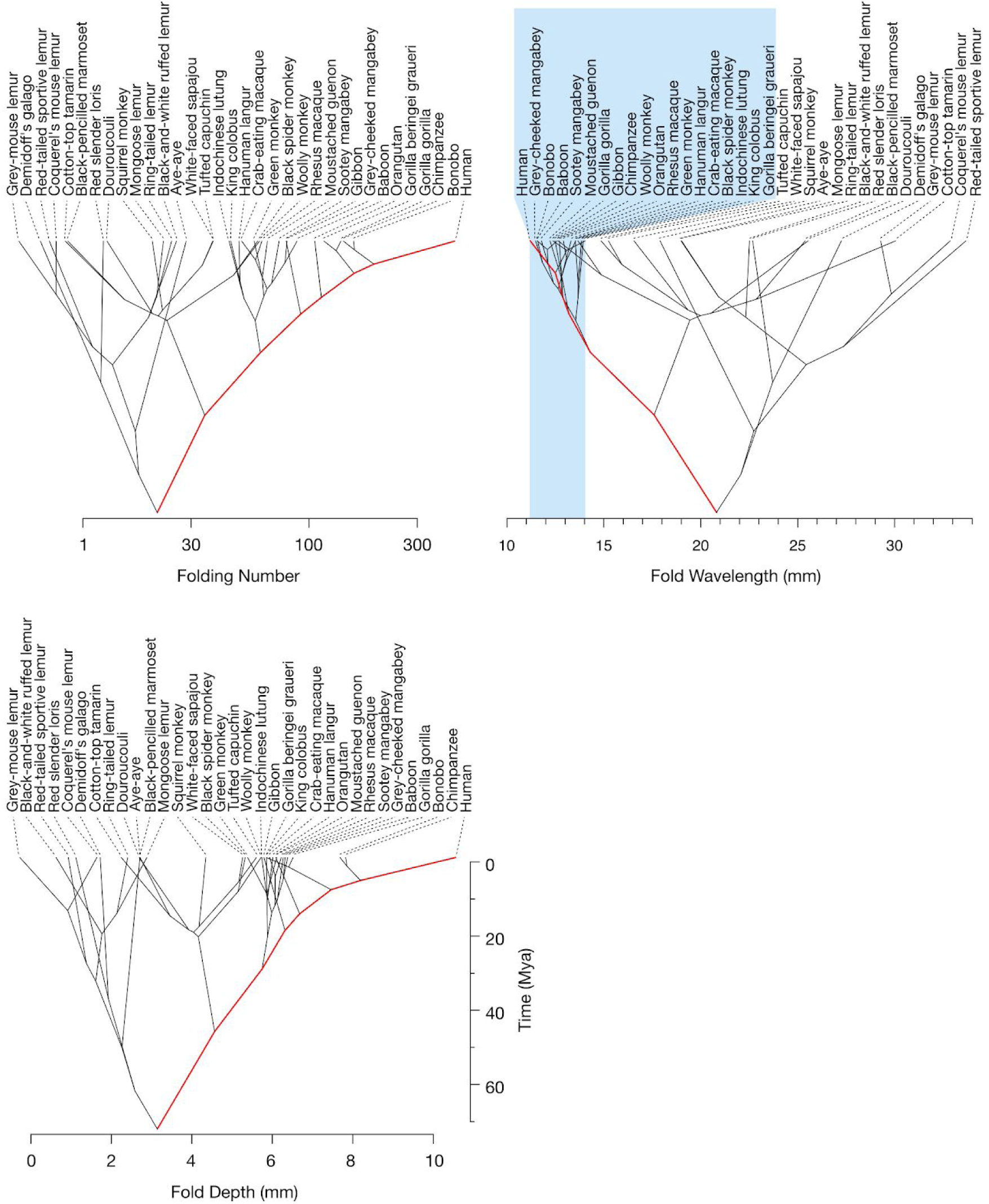
Phenograms of estimated ancestral neuroanatomical phenotypes. Phenograms provide an alternative visualisation of ancestral phenotype estimations. The value of each phenotype is represented in the x-axis against time in the y-axis. The tree nodes and tips are displaced to their estimated phenotype versus time positions. The estimation of phenotypic evolution along the branch leading from the common ancestor to humans is highlighted in red. The light-blue region in the phenogram for fold wavelength highlights the group of large-brain primate species whose fold wavelength ranges between 11 and 14 mm.

## Discussion

The study of the evolution of the primate brain should allow us to better understand the origin of our own cognition. It should also provide information on the sources of normal and pathological variability of human neuroanatomy — a major challenge for neuroscience today (Zilles and Amunts 2013). In a similar way as the analyses of genomes for multiple species allow us to detect highly conserved or rapidly evolving regions, an analysis of neuroanatomical evolution and conservation should allow us to detect the traces of evolution in different brain systems and regions. It should also allow us to evaluate the degree of phenotypic conservation across species, providing a framework to better understand natural variability, and to distinguish it from pathological variability.

It is difficult, however, to access and to analyse comparative primate brain data, by contrast to genetics where large open access databases such as GenBank (Benson et al 2017) provide information on thousands of different species. The series of reports by Stephan and Frahm (Stephan et al 1981, 1991, Frahm et al 1982) on regional brain volumes have been an important reference for the field, and their data tables have been used by many comparative brain analyses through the years. Primate brain MRI data is on the contrary only available for a few selected species. Two important resources are the PRIMate Data Exchange Initiative (PRIME-DE, Milham et al 2018, http://fcon_1000.projects.nitrc.org/indi/indiPRIME.html) and the National Chimpanzee Brain Resource (http://www.chimpanzeebrain.org). PRIME-DE shares open MRI data mostly for rhesus and crab-eating macaques, and NCBR shares open MRI data for several chimpanzee brains plus 9 other primate species (squirrel monkey, capuchin monkey, rhesus macaque, sooty mangabey, baboon, gibbon, orangutan, gorilla, and bonobo. Access to additional chimpanzee MRI data is available upon request). Another notable open data resource is the Macaque Neurodevelopmental Data project (Young et al 2017) which shares open longitudinal MRI data for 32 rhesus macaques. Although large MRI data samples have been acquired for other species by several groups (for example, Phillips and Sherwood 2008, Fears et al 2011, Love et al 2016), their access policy is less clear.

Here, we provide open access to a collection of 66 brain MRI datasets from 34 different primate species. These MRIs can be directly visualised and annotated in BrainBox using just a Web browser. In addition to indexing some of the data already online, we have scanned and made available 31 primate brain MRIs from 29 different species (23 species not previously available online), most of them with an isotropic resolution of 300 microns. We used Zenodo (https://zenodo.org) for storing the data. By using Zenodo each dataset is assigned a persistent identifier (digital object identifier, DOI), which facilitates data citation, tracking authorship and provenance. Other researchers willing to share their MRI data could similarly store it in Zenodo and index the URL in BrainBox. This would enable the decentralised creation of a community-curated collection of primate MRI data (the complete process of uploading the data and indexing it in BrainBox should not take more than 15 minutes). Using BrainBox, we were able to manually segment our MRI data, and to create topologically correct 3D surface reconstructions. The scripts necessary to programmatically download all our data and reproduce our statistical analyses are available on GitHub (https://github.com/neuroanatomy/34primates). Our aim is to make the data easily accessible to facilitate collaborative projects, reproducibility, and to encourage neuroscientists and citizen scientists to participate in advancing our understanding of primate brain diversity and evolution.

We have used this collection to study the variation and evolution of neocortical folding in primates. Previous comparative analyses of primate brain folding (for example, Prothero and Sundsten 1984, Zilles et al 1988, Rilling and Insel 1999, Semendeferi et al 2002, Lewitus et al 2014) have relied on 2-D measurements of gyrification indices. Using 3D meshes enables an extended set of neuroanatomical analyses to be performed, such as shape analyses, spectral analyses, surface-based alignment, among others. Here, we used surface-based maps of mean curvature to measure total folding length. Folding length varied from less than 4 cm in the grey-mouse lemur to up to 16 m in humans. We derived approximations of the average fold wavelength and fold depth. A well-known problem of the classical gyrification index (GI, Zilles et al 1988, 1989) is its difficulty to distinguish a brain with a few deep folds from one with a profusion of shallow folds. Zilles’s GI is computed for a coronal brain slice as the ratio between the pial contour and the contour of a hypothetical lissencephalic version of the brain. In 3D, Zilles’s GI is often approximated as the ratio between the neocortical surface and the surface of its convex hull, which exhibits again the same issue. Spectral analyses of brain folding offer a solution to the problem, however, their interpretation is not often trivial or intuitive. Our fold wavelength and fold depth estimations combine the measurement of cerebral surface area with the measurement of folding length to provide an intuitive decomposition of brain folding, free from the problem of GI-like estimations. Given two cerebra with the same surface area, similar convex hull (i.e., similar GI), but different number of folds, the one with the largest number of folds will also have a larger folding length, and in consequence a smaller fold wavelength and fold depth than the other.

We observed an interesting, non-linear relationship between fold wavelength and fold depth with cerebral volume. The fold wavelength and fold depth estimations were conceived with highly folded brains in mind (such as those of humans or other *Hominoidea*). In smaller, smoother brains, such as those of some *Strepsirrhini* and *Platyrrhini* primates, there are only 1 or 2 folds within each hemisphere, and it is not clear sometimes what a “gyrus” would be. In these cases, our fold wavelength estimations give estimates of about 3 cm, which corresponds more or less to the size of a complete hemisphere (as if the complete hemisphere were a single fold). As soon as the number of sulci increased, we observed that fold wavelength decreased and stabilised at a value of about 12 mm (± 20%) across different primate groups, and despite cerebral volumes ranging from ∼50cm^3^ (crab-eating macaque) to 1000 cm^3^ (humans) – a 20-fold variation. This stability in fold wavelength is in agreement with mechanical theories of neocortical folding (Toro and Burnod 2005, Toro 2012, Tallinen et al 2014, 2016, Foubet et al. 2018, Heuer and Toro 2019) which predict that fold wavelength should depend on the bending stiffness of the neocortex, strongly determined by cortical thickness. Indeed, the thickness of the neocortex changes very little across mammalian species (Mota and Herculano-Houzel 2015). In our sample, the cortical thickness of the small Demidoff’s galago was ∼1.5 mm (similar to that of a mouse), and ∼2.5 mm in humans (see also Fischl and Dale 2000). Cortical thickness, and consequently cortical bending stiffness should be relatively stable across primate species, leading to the stable fold wavelength that we observed in our data. The fast initial decrease in fold wavelength should be due to the emergence of new folds as the neocortex expands. Once neocortices are fully folded, the following slow decrease in fold wavelength could be related to frequency doubling – the formation of folds within folds – as observed in swelling gel experiments (Mora and Boudaoud, 2006). Neocortical mechanics could lead to the formation of stable neuroanatomical modules – the folds – which could then become the basis for the adaptation and selection of advantageous cytoarchitectonic, connective and functional organisations, a kind of mechanical canalisation process (Waddington 1942, Müller 2007, Foubet et al 2018, Heuer and Toro 2019). A future analysis of fold wavelength and thickness, potentially local instead of only global, should allow us to better understand this relationship across and within species.

Our phylogenetic comparative analyses suggested that random phenotypic change may be an important driving force in the evolution of primate neocortical folding. After fitting several alternative evolutionary models, the Brownian Motion (BM) model captured better the variability in the data than the Ornstein-Uhlenbeck (OU) and Early-Burst (EB) models (2^nd^ and 3^rd^ best ones). The difference in fitting quality was not enough, however, to outrule the OU and EB models. Future analyses with larger samples should allow us to progress further in this respect. While the BM model supposes that phenotypic variation along the phylogenetic tree is random, the OU and EB models suppose the presence of advantageous phenotypes which drive evolution. It is important to note that the BM model is not incompatible with adaptive evolution (Nunn 2011). The driver of the random changes can still be natural selection, but with changes in the selective regime independent of previous changes and more common along longer branches (probably due to rapidly changing environmental conditions). In all cases, the importance of phylogenetic relationships was strong and highly statistically significant: a star phylogenetic model – one where all species are considered to be independent – had a substantially less good fit to the data than the top 3 models.

Based on the BM model, the common ancestor of all primates, 74 million years ago, may have had a cerebrum similar to that of a small lemur: with a surface area of 50 cm^2^, a volume of 12 cm^3^, an absolute gyrification index of 2, a folding length of 37 cm, and about 25 folds, of an average wavelength of 20 mm and with a depth of about 3 mm, that is, not very different from that of a mongoose lemur or an aye-aye. Our estimation of global gyrification (AbsGI = 2.1, 95% CI from 1.3 to 2.8) is not much higher than that provided by the previous phylogenetic comparative analysis of Lewitus et al (2014), which gave a GI=1.41 for the common ancestor of primates (AbsGI is also expected to be higher than GI). The increase in volume and gyrification observed in the large *Catarrhini* (the group containing humans but also macaques) may have started about 40 million years ago, but probably only about 7 million years ago (about the time of the last common ancestor of humans and chimpanzees) in the branch leading to *Cebidae*, such as the white-faced sapajou or the tufted capuchin. Lissencephaly, as observed in *Platyrrhini*, such as the cotton-top tamarin and the black-pencilled marmoset, may have evolved from a gyrencephalic ancestor about 20 million years ago.

Lewitus et al (2014) and more recently Lewitus et al (2016) have suggested that the process leading to gyrencephaly may have emerged at least twice during mammalian evolution. As an example of such process they cite the results of Reillo et al (2010) or more recently De Juan Romero et al (2015). These studies suggest that gyri are produced by local bulging due to a genetically programmed increase in neurogenesis. Instead of explaining the evolutionary gain or loss of folding by a complex readjustment of the genetic patterning of the neocortex, it seems to us that mechanical theories provide a more parsimonious explanation for our data: neocortical folding would appear and disappear as soon as neocortical growth relative to the growth of the white matter substrate goes beyond or under the mechanical buckling threshold. The highly conserved fold wavelength that we observed would simply reflect a similar neocortical stiffness across species instead of a more complex genetic patterning process appearing and disappearing through the ages. Within the context of mechanically produced folding, genetics would have a more subtle role, providing a meta-level of regulation and selection of structures which appear by physical necessity instead of a detailed prescription of each fold. Some small insects are able to stand on top of the water in a pond, an ability that larger insects do not exhibit. It seems more parsimonious to explain this through the water’s surface tension than by a complex cascade of genetic processes leading to the ability of very different species of larger insects to sink.

## Acknowledgements

We thank Marion Fouquet and Nicolas Traut for their help with the quality control and selection of the human data from ABIDE. We thank Helen D’Arcueil, Alex de Crespigny, Emmanuel Gilissen and Chet Sherwood for allowing us to make accessible their MRI data in the Brain Catalogue. We thank Spencer Arbuckle and Andrew Pruszynski for sharing their macaque data with us and making it openly available. Data acquisition was funded by the MNHN programme e-Museum. The development of BrainBox was supported by the Wellcome Trust through the Open Science Prize. KH was supported by the Max Planck International Research Network on Aging and the Max Planck Institute for Human Cognitive and Brain Sciences. OFG was supported by NWO VIDI grant 864-13-012.

